# Stochasticity and negative feedback lead to pulsed dynamics and distinct gene activity patterns

**DOI:** 10.1101/011734

**Authors:** Samuel Zambrano, Marco E. Bianchi, Alessandra Agresti, Nacho Molina

**Affiliations:** San Raffaele University, Via Olgettina 58, 20132 Milan, Italy.; San Raffaele Scientific Institute, Division of Genetics and Cell Biology, Via Olgettina 60, 20132 Milan, Italy.; SynthSys Center, University of Edinburgh, Mayfield Road, EH9 3JD. United Kingdom.

## Abstract

Gene expression is an inherently stochastic process that depends on the structure of the biochemical regulatory network in which the gene is embedded. Here we study the interplay between stochastic gene switching and the widespread negative feedback regulatory loop. Using a simple hybrid model, we find that stochasticity in gene switching by itself can induce pulses in the system. Furthermore, we find that our simple model is able to reproduce both exponential and peaked distributions of gene active and inactive times similar to those that have been observed experimentally. Our hybrid modelling approach also allows us to link these patterns to the dynamics of the system for each gene state.

---

Cells need to provide an adapted response to external stimuli, which requires the production of the adequate proteins following different temporal patterns. This is achieved through biochemical networks in which a stimulus triggers a cascade of reactions that eventually lead to the activation of transcription factors, proteins that activate or repress the expression of specific gene sets. Thus, the temporal regulation of gene activity will be determined by the structure of the network in which the gene is embedded [1]. A common regulatory structure is the negative feedback loop, in which a transcription factor activates the production of a protein that contributes to its own inhibition. This motif regulates the activity of important transcription factors such as NF-*κ*B [2] and p53 [3] and has been shown to give rise to pulses in the concentration of the proteins of the network (see e.g. [4, 5]) as predicted by mathematical models [6]. The role of this pulsed dynamics is not fully understood though: theoretical studies suggest that they can give rise to a more reliable protein production [7] while experiments show that oscillatory dynamics can determine the cell fate [8].

On the other hand, gene expression is an intrinsically noisy process [9] and models in which the gene state switches randomly between active and inactive states are able to fit well experimental data [10]. By contrast, some genes show peaked distributions of active/inactive times [11–13] that deviate from the exponential distributions obtained with simple random models. These distributions can be obtained using multi-step models mirroring the multiple steps of gene activation [14], but they could also arise from the interplay between the stochastic gene activity and the structure of the regulatory network in which the gene is embedded. Some insights about this interplay have been gained by showing the emergence of oscillations when a gene is an autorepressor [15], of noise-enhanced persistence of biochemical species [16] and how stochasticity dephases genetic oscillators [17]. A major obstacle in this context is the difficulty of providing analytical insights on the nonlinear stochastic systems involved (often modelling species with very low copy numbers). For this reason, we are far from having a complete picture of the type of regulation that emerges from such interaction.

In this Letter we describe the dynamics emerging from the interaction of stochastic gene switching and the widespread negative feedback in a simple network using a hybrid modelling approach, in which only the gene activity is modelled as a stochastic process. Our hybrid simulations show that stochastic gene switching is responsible for most of the dynamical variability and induces pulsed dynamics in the system, though the deterministic model predicts a steady state. We show that even in this simple biochemical network distinct dynamical patterns of gene activity can arise, and how hybrid modelling allows us to gain analytical insight on their origin. We discuss the implications of our results in the end of this Letter.

The model considered here is shown in Fig. 1(a). This model adds a layer of regulation to the one proposed in Ref. [7] and is a simplified version of the biochemical network of NF-*κ*B [18]. It is formed by a gene that can be active G or inactive 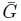 and an *activator A* that can activate the gene, similarly to NF-*κ*B [18]. When the gene is active, the *inhibitor* protein *I* is produced and provides the negative feedback by both contributing to the gene's inactivation and by forming a complex with *A* that cannot activate the gene any longer. In what follows we use the same letters both for the names of the biochemical species and for their copy numbers. For the sake of simplicity we consider that we have only one copy of the gene, so 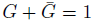 and that the total amount of *A* (free and bound to *I*) is constant and equal to *A*_tot_, as for NF-*κ*B [18]. Finally we assume that the inhibitor undergoes degradation both in the free and in the complex form.

**Figure 1:**
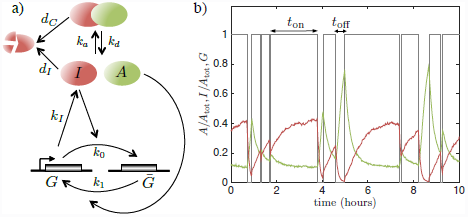
Dynamics of a network with stochastic gene switching and a negative feedback loop, (a) Model diagram showing the activator *A*, the inhibitor, *I*, and the gene that can be active *G* or inactive 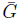. (b) A stochastic simulation of a trajectory of our system showing the free activator fraction *A/A*_tot_ (green), the gene state *G* (grey) and the relative amount of inhibitor *I/A*_tot_ (red) displaying pulses. For the gene state, an active time (*t*_on_) and an inactive time (*t*_off_) are displayed. Simulations were performed with parameters *d*_C_ = 1.5 · 10^−4^*s*^−1^, *k*_a_ = 5.1 · 10^−6^*s*^−1^, *k*_*d*_ = 1.4 · 10^−3^*s*^−1^, *d*_I_ = 2.8 · 10^−4^*s*^−1^, *k*_I_ = 11.2 *s*^−1^, *k*_0_ = 1.4 · 10^−7^ *s*^−1^, *k*_1_ = 5.8 · 10^−7^*s*^−1.^

In Figure 1(b) we show a stochastic trajectory of the system obtained using the Gillespie algorithm [19]. It exhibits pulses of the free activator *A* and the rest of variables, as observed in different networks with negative feedbacks [6]. The parameters used were obtained from Ref. [18] and adjusted to obtain *O*(10^4^) copies of each protein and inhibitor half-life and activator intra-peak timing of the order of one hour, the typical timescale of these biological oscillators [6]. Pulses are spiky as in models of oscillations of NF-*κ*B [6] although we have found that in stochastic simulations of slightly more complex biochemical networks (e.g. [18]) a wider variety of pulses arises.

Another traditional modelling approach for biochemical networks is to use ordinary differential equations derived from mass-action kinetics [20], but this is inadequate when species with low copy numbers are present (in our case, *G*). High copy numbers is also a necessary condition to approximate the dynamics of the system by using a Langevin equation, although exact results can only be obtained for linear biochemical networks [21]. For these reasons, in this kind of systems the so-called hybrid simulations [22] in which part of the reactions are modeled by a Langevin equation [23] while the rest are modeled as stochastic processes, are increasingly popular. This procedure gives simulations that mimic the behaviour of the fully stochastic system [22] and significantly reduce the computation time.

Inspired by these algorithms we study our system through a *simplified hybrid* model in which the only species modelled stochastically is the gene state G. This type of modelling has indeed already been used to simulate complex models of cell signalling, see e.g. [24–26]. But most importantly, this kind of modelling allows us to isolate and identify in a precise way the role of stochastic gene activity in the dynamics.

Using our hybrid approach we model the evolution of *A* and *I* as:

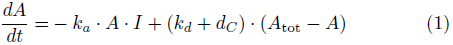

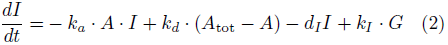

This nonlinear dynamical system is driven by the stochastic process of gene switching:

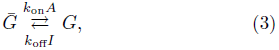

where the switching rates will depend on time through the variables *A* and *I.* Thus, for the gene state *G* we can write down the chemical master equations:

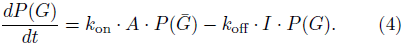

Numerical simulations of this hybrid model are performed using a deterministic integrator for Eqs. 1 and 2 and switching the value of *G* between *G* = 0 and *G* = 1 following the Gillespie algorithm, as prescribed in Ref. [27]. For a fully deterministic simulation of the model using mass-action kinetics it would be enough to add to equations 1 and 2 the following equation,

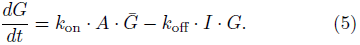

which in the equilibrium 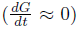 leads to a Michaelis-Menten like equation for *G* [1].

In Figure 2(a) we show the evolution in time of the free activator *A* obtained for fully stochastic, hybrid and deterministic simulations. We can observe that the pulses obtained for the fully stochastic simulations and the simplified hybrid simulations are very similar. Interestingly, we observe that the deterministic simulations lead to the convergence of the system to a steady state. From this we conclude that stochastic gene switching by itself can induce pulses in the network. This is another example of how stochasticity can induce pulses in contexts where deterministic models predict steady states, as observed in models of population dynamics [28] and excitable systems [29].

**Figure 2:**
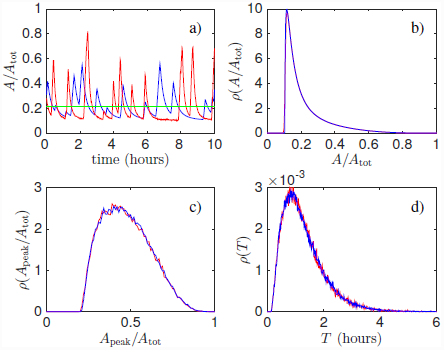
Stochastic gene switching is responsible for most of the dynamic variability in the network. (a) Evolution in time of the free activator A using fully stochastic (red) and simplified hybrid (blue) simulations. Both display similar pulses while the fully deterministic simulation (green) remains at a stable fixed point. (b) Distribution of the values of A for both the hybrid (blue) and fully stochastic (red) simulations of the biochemical network. (d) Distribution of the amplitudes and (e) the periods of the pulses for the hybrid (blue) and fully stochastic (red) model. The distribution of the activator and the pulse parameters are very similar for the hybrid and the fully stochastic system.

Simulations show that stochastic gene activity is responsible for most of the variability of the system. In Fig. 2(b) we show the distributions of *A*, represented by their probability density *ρ*, for stochastic and hybrid simulations, which are nearly indistinguishable. On the other hand, by using a simple peak detection algorithm we can detect the timing between two consecutive peaks *T* and their amplitude *A*_peak_. For these calculations we consider only peaks of at least 10% of *A*_tot_, the order of magnitude that can be detected in experiments of activator dynamics such as NF-*κ*B [5]. The distributions of these magnitudes are shown in Fig. 2(c) and (d) respectively, and are again nearly indistinguishable: this confirms the crucial role of stochastic gene activity in the dynamic variability of the system and the ability of hybrid model to mimic the fully stochastic simulations (in drastically shorter computation times).

Hybrid modelling also allows us to understand the pulsed dynamics of our network in terms of the the null-clines of the system given by Eqs. 1 and 2. There are three of such nullclines: the one that we obtain by setting 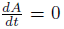, *I* = *f_A_*(*A*), and the two nullclines that we obtain by setting 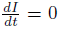 for *G* = 0 and *G* = 1, denoted *I* = *f*_*I*,0_(*A*) and *I* = *f*_*I*,1_(*A*) respectively. The nullclines and a trajectory for our hybrid model are depicted in Fig. 3. It is easy to see that irrespectively of the parameter values *I* = *f*_*A*_(*A*) intersects in exactly one point, 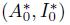, with *I* = *f*_*I*,0_(*A*) and in exactly one point, 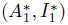, with *I* = *f*_*I*,1_(*A*). Furthermore, an analysis on the direction of the flow determined for each gene state shows that these two fixed points are necessarily stable. Thus, for this simple biochemical network the pulses can be understood as a series of jumps between the fixed points obtained for *G* = 0 and for *G* = 1.

**Figure 3:**
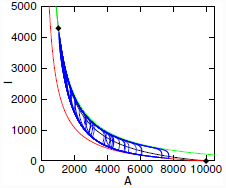
The three nullclines that can be associated to the hybrid model: the nullcline for 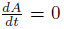 (black line) and the nullclines for 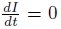 with *G* = 1 (green line) and *G* = 0, respectively. A trajectory of the hybrid model is also depicted (blue line). There are two intersection between these three curves (black diamonds); a simple graphical analysis shows that these points are stable. Trajectories of the hybrid model jump between these two fixed points.

Experimental studies in which gene expression is monitored in real time have shown that gene active and inactive times can either be exponentially distributed or be described by peaked Gamma-like distributions [11–13]]. These distributions are important signatures of the underlying stochastic process that drives gene activity. To explore the gene active and inactive distributions that our simple model can generate, we study the dynamics of the system in parameter space by varying randomly each of the parameters within one order of magnitude from the values used in our previous simulations. For each parameter values, we simulated the dynamics and obtained the histograms of the active and inactive times (*t*_on_ and *t*_off_, see Fig. 1(b)). We grouped them in ten clusters according to their coefficient of variation (CV) i.e. the standard deviation divided by the mean. We found that *t*_on_ and *t*_off_ can be distributed following quite distinct patterns: we observed distribution with shapes that range from an exponential-like shape, with a global maximum at zero and high CV, to a Gamma-like shape, with a global maximum close to the mean and low CV (see Fig. 4(a) and (b)). Our simple system is then able to recapitulate the experimentally observed distributions. In particular our calculations show that a negative feedback can give rise to peaked distributions and thus can be an alternative to the multi-step process models proposed to explain the distributions observed in experiments [11, 13].

**Figure 4:**
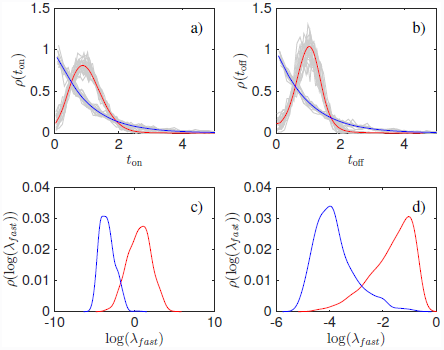
Distinct gene activity patterns are reproduced with different model parameters. (a) Distributions of the gene active times *t*_on_ and (b) inactive times *t*_off_ with the highest and the lowest CVs, average in blue and red, respectively. (c) Distributions of the eigenvalues with bigger absolute value *λ*_*fast*_ of the fixed point 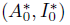 (obtained when the gene is inactive) and (d) of the fixed point 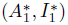 (obtained when the gene is active), producing high-CV distributions (blue) and low-CV distributions (red).

Using the hybrid modelling we can investigate the origin of these patterns. For the sake of simplicity we focus on the distribution of the active times (*t*_on_). For our system, if at time *t* = 0 the system is in the state *G*, the probability of remaining in the state *G* can be expressed in terms of the conditional probability *P*_*G*_(*t*|*V*_0_) given the initial condition *V*_0_ = (*A*_0_, *I*_0_) and the probability 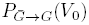 of finding the system at *V*_0_ at the initial gene transition 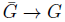 as:

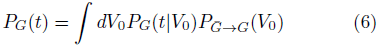

where the conditional probability is:

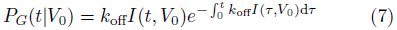

Notice that if the overall switching rate is constant (*k*_off_*I*(*t*, *V*_0_) = *cte*), as in the random switching model [10], an exponential probability distribution for the gene inactive time will be recovered. Instead, we find that in some cases the probability distribution is nonexponential. In this situation we would expect that the conditional distributions *P*_*G*_(*t*|*V*_0_) that contribute the most to the integral in Eq. 6 should have a relative maximum at *t_max_* ≠ 0. Such relative maximum would satisfy the following equation,

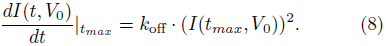

Solving this equation requires the explicit form of *I*(*t*, *V*_0_). However we know that 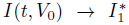 and 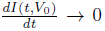 as *t* → ∞. Hence, equation 8 is the equation of the intersection of a monotonically decreasing function 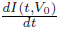 (trajectories are always below the nullcline *I* = *f*_*I*,1_(*A*) and 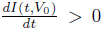, see Fig. 3) and a monotonically increasing function *k*_off_ · (*I*(*t,V*_0_))^2^ (because the amount of inhibitor grows when *G* = 1, see again Fig. 3) at the point *t* = *t*_*max*_. Our previous nullclines analysis shows that 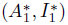 is a stable fixed point with negative eigenvalues with absolute values *λ*_*fast*_ > *λ*_*slow*_. Considering all this, we can roughly approximate 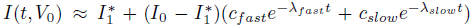 where *c*_*fast*_ > 0, *c*_*slow*_ > 0 and *c*_*fast*_ + *c*_*slow*_ = 1. For short times (compared with 1/*λ*_*slow*_) the derivative 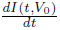 scales with *λ*_*fast*_ and hence the crossing determined by Eq. 8 will only be possible if *λ*_*fast*_ is sufficiently big (and *k*_off_ is sufficiently small). It is easy to show that the same argument leads to an equivalent result for the distribution of the inactive times (*t*_off_) where *λ*_*fast*_ is the corresponding fast eigenvalue at the fixed point with *G* = 0.

We provide a numerical confirmation of the validity of this argument in Figs. 4(c) and (d), where we show that the values of *λ*_*fast*_ for the corresponding fixed points (for *G* = 0 and *G* = 1) are able to discriminate between the two different gene activity patterns, since the larger the eigenvalue is, the more likely is that *t*_off_ (and *t*_on_) are exponentially distributed. Thus, the use of hybrid modelling allows to gain insights on the origin of the patterns of gene activity observed.

Pulsed dynamics is widespread in genetic circuits [30]. Our simple model shows that the interplay of negative feedbacks and stochastic gene switching gives rise to pulsed dynamics even if the fully deterministic simulations predict convergence to a steady state. Furthermore, we have found that in spite of the simplicity of the dynamics arising, our network can display different gene activity patterns. The negative feedback plays a key role in this fine temporal control: without it, the dynamics of gene activation would be purely random. Our results imply that in experiments in which the gene activity patterns are found to be peaked [11–13]] a negative feedback loop might be at work. We think that the increasing availability of experimental data will allow to delineate the contribution to the gene activity dynamics by both the multi-step sequential stochastic process of gene activation and the constraints imposed by the structure of the regulatory biochemical network. As in our work, the use of hybrid modelling can help to provide further analytical insights.

SZ has been partially supported by the Intra-European Fellowships for career development-2011-298447NonLinKB. NM has been fully supported by the Chancellor’s Fellowships granted by the University of Edinburgh.

## References

[1] U. Alon, An Introduction to Systems Biology (CRC Press, Boca Raton, FL (USA), 2007).

[2] A. Hoffmann, A. Levchencko, M. Scott, and D. Baltimore, Science 298, 1241 (2002).

[3] N. Geva-Zatorsky, N. Rosenfeld, S. Itzkovitz, R. Milo, A. Sigal, E. Dekel, T. Yarnitsky, P. Pollack, Y. Liron, Z. Kam, et al., Molecular Systems Biology 2, E1 (2006).

[4] D. Nelson, A. E. Ihekwaba, M. Elliott, J. Johnson, C. A. Gibney, B. Foreman, G. Nelson, V. See, C. Horton, D. Spiller, et al., Science 306, 704 (2004).

[5] S. Zambrano, M. E. Bianchi, and A. Agresti, PLoS ONE 9, e90104 (2014).

[6] G. Tiana, S. Krishna, S. Pigolotti, M. H. Jensen, and K. Sneppen, Phys. Biol. 4, R1 (2007).

[7] F. Tostevin, W. de Ronde, and P. R. ten Wolde, Phys. Rev. Lett. 108, 108104 (2012).

[8] J. E. Purvis, K. W. K. C. Mock, E. Batchelor, A. Loewer, and G. Lahav, Science 336, 1440 (2012).

[9] M. B. Elowitz, A. J. Levine, E. D. Siggia, and P. S. Swain, Science 297, 1183 (2002).

[10] A. Raj, C. S. Peskin, D. Tranchina, D. Y. Vargas, and S. Tyagi, PLoS Biology 4, e309 (2006).

[11] D. M. Suter, N. Molina, D. Gatfield, K. Schneider, and U. Schibler, Science 332, 472 (2011).

[12] C. V. Harper, B. Finkenstädt, D. J. Woodcock, S. Friedrichsen, S. Semprini, L. Ashall, D. G. Spiller, J. J. Mullins, D. A. Rand, J. R. E. Davis, et al., PLoS Biology 9, e1000607 (2011).

[13] N. Molina, D. M. Suter, R. Cannavo, B. Zoller, I. Gotic, and F. Naef, PNAS 110, 20563 (2013).

[14] A. Coulon, C. C. Chow, R. H. Singer, and D. R. Larson, Nature Rev. Gen. 14, 572 (2013).

[15] P. E. Morant, Q. Thommen, F. Lemaire, C. Vaandermoere, B. Parent, and M. Lefranc, Phys. Rev. Lett. 102, 068104 (2009).

[16] M. Assaf and B. Meerson, Phys. Rev. Lett. 100, 058105 (2008).

[17] D. A. Potoyan and P. G. Wolynes, Proc. Natl. Acad. Sci. USA 111, 2391 (2014).

[18] S. Zambrano, M. E. Bianchi, and A. Agresti, J. Theor. Biol. 347, 44 (2014).

[19] D. T. Gillespie, J. Phys. Chem. 81, 2340 (1977).

[20] V. Chellaboina, S. Bhat, M. Haddad, and D. Bernstein, Control Systems, IEEE 29, 60 (2009).

[21] P. B. Warren, S. Tanase-Nicola, and P. R. ten Wolde, J. Chem. Phys. 125, 144904 (2006).

[22] H. Salis and Y. Kaznessis, J. Chem. Phys. 122, 054103 (2005).

[23] D. T. Gillespie, Am. J. Phys 64, 1246 (1996).

[24] S. Tay, J. J. Hughey, T. K. Lee, T. Lipniacki, S. R. Quake, and M. W. Covert, Nature 466, 267 (2010).

[25] P. Paszek, S. Ryan, L. Ashall, K. Sillitoe, C. V. Harper, D. Spiller, D. A. Rand, and M. White, Proc. Natl. Acad. Sci. USA 107, 11644 (2010).

[26] J. R. Karr, J. C. Sanghvi, D. N. Macklin, M. V. Gutschow, J. M. Jacobs, B. Bolival, N. Assad-Garcia, J. I. Glass, and M. W. Covert, Cell 150, 389 (2012).

[27] T. Lipniacki, K. Puszynski, P. Paszek, A. Brasier, and M. Kimmel, BMC Bioinformatics 8, 366 (2007).

[28] A. J. McKane and T. J. Newman, Phys. Rev. Lett. 94, 218102 (2005).

[29] P. Rué and J. Garcia-Ojalvo, Mathematical biosciences 231, 90 (2011).

[30] J. H. Levine, Y. Lin, and M. B. Elowitz, Science 342, 1193 (2013).

